# Comparative genomics analysis reveals that antimicrobial activity in *Pseudomonas protegens* PBL3 is associated with gene clusters participating in multiple cellular functions

**DOI:** 10.64898/2026.02.12.705552

**Authors:** Shilu Dahal, Chia Sin Liew, Jean-Jack Riethoven, Clemencia M. Rojas

## Abstract

The environmental bacterium *Pseudomonas protegens* PBL3 has antagonistic activity against the bacterial pathogen *Burkholderia glumae,* the causal agent of the rice disease Bacterial Panicle Blight (BPB). The antimicrobial activity of the *P. protegens* PBL3 was found in the bacteria-free secreted fraction (secretome), but the specific molecules, as well as the genetic basis conferring activity, have not been identified. To elucidate the genes and biosynthetic pathways governing the antimicrobial activity of the *P. protegens* PBL3 secretome, we used comparative genomics, by leveraging six *Pseudomonas* spp. strains, with available genomic sequences and exhibiting contrasting antimicrobial activities against *B. glumae*. We hypothesized that *Pseudomonas* spp. strains with antimicrobial activity against *B. glumae* have conserved genes with *P. protegens* PBL3 and those genes are absent in the strains lacking activity. To test this hypothesis, we performed comparative genomics analysis across the six strains using two complementary approaches, anvi’o and progressiveMauve, and using *P. protegens* PBL3 as the reference genome. This analysis revealed 188 genes uniquely present in antimicrobial-producing strains. Seven of those genes were associated with biosynthetic gene clusters predicted to encode secondary metabolites; additional genes were grouped into twenty-five contiguous clusters with functions associated with secretion, signal transduction, regulation, transport/efflux, carbohydrate metabolism and one with additional uncharacterized function. Altogether, this study provided a complex and multi-functional network of candidate genes in antimicrobial-producing strains, suggesting that *P. protegens* PBL3 employs not only classical biosynthetic pathways but also integrated regulatory, metabolic, and export modules to synthesize and deploy antimicrobials.

**Importance:** Bacterial plant diseases can be controlled through biological control, a strategy wherein beneficial microorganisms known as biological control agents (BCAs) interfere with the biological activities of pathogens through several mechanisms. One of these mechanisms is antibiosis, by which antimicrobial molecules produced by the BCA prevent pathogen multiplication or actively kill the pathogen. While this mechanism of antibiosis has been widely recognized, the specific molecules associated with the antimicrobial activity are not always identified, given their diverse and complex chemical structures as well as the unique and intricate biosynthetic pathways. This study unraveled different pathways underlying the antimicrobial activity in the environmental bacterium *Pseudomonas protegens* PBL3 to advance the discovery of effective antimicrobials to control bacterial plant diseases.

## Introduction

Plant diseases can be effectively controlled by the biological activities of beneficial microorganisms in the ecosystem, an approach called biological control (Collinge et al., 2022). Biological control agents (BCAs) interfere with the activities of plant pathogens either indirectly, by inducing plant resistance or by competing for nutrients or space with pathogens, or directly, by antagonism through the production of antimicrobial molecules that kill or reduce the growth of pathogens (Conrath et al., 2015; Kohl et al., 2019; Pal & McSpadeden Gardener, 2006; Pieterse et al., 2014)

The genus *Pseudomonas* is known for its remarkable metabolic diversity, synthesizing a wide array of secondary metabolites with potent antimicrobial properties, which have resulted in several members of the genus to be registered as BCAs against pathogens causing diseases in diverse crops (Alattas et al., 2024; Gross & Loper, 2009; Mishra & Arora, 2018; Yang et al., 2025). Accordingly, our previous work uncovered the antimicrobial activity of *Pseudomonas protegens* PBL3 against *Burkholderia glumae*, the causal agent of the rice disease Bacterial Panicle Blight (BPB) (Fory et al., 2014; Ham et al., 2011; Ortega & Rojas, 2021; Ortega et al., 2020; Zhou, 2019). We also found that this antimicrobial activity was associated with molecules secreted by the bacteria (secretome) likely associated with putative secondary metabolites encoded by biosynthetic gene clusters (BGCs) predicted by the *P. protegens* PBL3 genomic sequence (Ortega et al., 2020). To gain insight into the function of *P. protegens* PBL3 secretome antimicrobial activity, we tested 8 of the genome-predicted compounds against *B. glumae* using commercially available analogs, and found that only Pyoverdine, Pyoluteorin, and 2,4-Diacetylphloroglucinol (2,4-DAPG) inhibited the growth of *B. glumae* at high concentrations (Dahal et al., 2024). These molecules, however, did not fully explain the observed antimicrobial activity of the *P. protegens* PBL3 secretome as quantification of their endogenous concentrations by reversed-phase liquid chromatography with tandem mass spectrometry (RPLC-MS/MS), using the commercial analogs as standards, revealed that their endogenous concentrations were below the concentrations needed for activity (Dahal et al., 2024). The latter finding suggests that either the antimicrobial activity of the *P. protegens* PBL3 secretome is associated with the combination of known antimicrobial molecules, other uncharacterized secondary metabolites encoded by these known BGCs, or by novel structurally and chemically diverse molecules encoded in other genomic regions.

The identification of naturally-occurring antimicrobials is challenging due to their immense structural diversity, their production under specific growth/environmental conditions, and technical limitations with bioanalytical approaches and compound identification (Krug & Muller, 2014; Nguyen et al., 2024; Santamaria et al., 2022; van der Hooft et al., 2020). The alternative approach to facilitate the identification of genomic regions associated with antimicrobial activity is through comparative genomics (Calderón et al., 2015; Chung et al., 2021; Panter et al., 2021; Saati-Santamaria et al., 2022).

Here, we applied a comparative genomics framework to identify genes and biosynthetic pathways underlying the antimicrobial activity of *P. protegens* PBL3 against *B. glumae*. To facilitate this approach, we selected six closely related *Pseudomonas* spp. strains with available genome sequences and differential antimicrobial activity against *B. glumae.* This differential antimicrobial activity led us to hypothesize that *Pseudomonas* spp. strains with antimicrobial activity against *B. glumae* have genomic regions in common with *P. protegens* PBL3, that are absent in *Pseudomonas* spp. strains without such antimicrobial activity. To test this hypothesis, we conducted a comparative genomic analysis among those strains and using the *P. protegens* PBL3 as reference genome to identify those genomic regions uniquely present in the *P. protegens* PBL3 genome. This approach resulted in the identification of 188 annotated non-redundant putative genes. These genes span biosynthetic enzymes, metabolism, transporters, secretion systems, and regulatory components, suggesting that antimicrobial activity in *P. protegens* PBL3 is not only related to antimicrobial biosynthetic pathways but by an integrated, multi-functional genetic network. This integrated architecture supports a model in which antimicrobial activity in *P. protegens* PBL3 arises from the combined action of genes associated with metabolite production, regulation, and efficient export or delivery mechanisms.

## Materials and Methods

### Bacterial strains

Bacterial strains used in this study are listed in Table 1. *Burkholderia glumae* UAPB13 was grown in the King’s B (KB) media (King et al., 1954) and all the *Pseudomonas* strains were grown on Luria Bertani (LB) medium (Bertani, 1951). Genomes of the *Pseudomonas* strains were obtained from NCBI and JGI Integrated Microbial Genomes & Microbiomes (IMG/M) (Chen et al., 2022).

**Table 1.** Bacterial strains used in this study.

| Strains | Source | NCBI GenBank or JGI<br>IMG Taxon ID |
| --- | --- | --- |
| *Burkholderia glumae* UAPB13 | Rice panicle | GCA_024611525.1 |
| *Pseudomonas protegens* PBL3 | Soil sample from wheat fields | GCA_016903635.1 |
| *Pseudomonas protegens* 3295 | Soil sample from sweet sorghum fields | 2818991464 |
| *Pseudomonas fluorescens* 513 | Roots from pearl forage millet | 2834028612 |
| *Pseudomonas fluorescens* 1550 | Rhizosphere from grain sorghum fields | 2923586266 |
| *Pseudomonas fluorescens* 4488 | Rhizosphere from grain sorghum fields | 2904518522 |
| *Pseudomonas fluorescens* 1098 | Rhizosphere from energy sorghum | 2919063839 |
| *Pseudomonas putida* 4512 | Roots from false purple brome | 2919228037 |

### Secretome preparation and *in vitro* antimicrobial assay

Cell-free secretions (secretome) were harvested from *P. protegens* PBL3, *P. fluorescens* 513, *P. fluorescens* 1550, *P. protegens* 3295, *P. fluorescens* 1098, *P. fluorescens* 4488, and *P. putida* 4512 for an antimicrobial assay against *B. glumae*, following previously described procedures (Dahal et al., 2024).

### Genome quality assessment and improvement

The quality of genomes was evaluated using QUAST (Quality Assessment Tool for Genome Assemblies) v5.0.2 (Gurevich et al., 2013). The draft genomes of the six *Pseudomonas* strains (*P. fluorescens* 513, *P. fluorescens* 1550, *P. protegens* 3295, *P. fluorescens* 1098, *P. fluorescens* 4488, and *P. putida* 4512) were enhanced by RagTag v2.1.0 (Alonge et al., 2022) and using the *P. protegens* PBL3 genome as the reference to order and orient assembled contigs into longer sequences (scaffolds) by inserting gaps where the sequence is unknown.

### Identification and comparison of biosynthetic gene clusters (BGCs) across *Pseudomonas* spp. strains

Secondary metabolite BGCs were predicted using Antibiotics & Secondary Metabolite Analysis Shell (antiSMASH) v8.0.4 (Blin et al., 2025) for all seven *Pseudomonas* genomes analyzed in this study. Each genome was processed with default parameters, enabling detection of secondary metabolite classes. BGC annotations generated for each strain were compiled into a presence absence matrix to facilitate cross-strain comparison in reference to *P. protegens* PBL3.

### Comparative genomic analysis

Comparative genomic analyses were performed using complementary pangenome and whole-genome alignment approaches to identify genomic regions associated with antimicrobial activity. Comparative pan-genomic analyses were performed using anvi’o (an analysis and visualization platform for ‘omics data) v8.0 (Eren et al., 2021) to estimate relationships among the genomes based on gene clusters (Delmont & Eren, 2018). For the pangenomics workflow, annotated contigs-dbs were first created for the seven genomes, and then used to generate a genomes-storage-db. This genome storage database was then used to run the pangenomic analysis using the command anvi-pan-genome. The contigs-dbs were annotated with the NCBI’s 2020 release of COGs (COG20) and the KEGG databases v2023-09-22. Genomic regions of interest were defined as those present in the genomes of bacterial strains with antimicrobial activity and lacking in the genomes of bacterial strains without antimicrobial activity. To extract these regions, a matrix of every gene cluster and its frequency of occurrence in each genome was obtained from the pangenomic profile (Shaiber et al., 2020).

To complement the pan-genomic analysis and assess large-scale genome organization and synteny, whole-genome alignments were performed using progressiveMauve v2.4.1 with default parameters (Darling et al., 2004; Darling et al., 2010), to identify conserved, orthologous regions among all or subsets of the input genomes that are free from genome rearrangements (Locally Collinear Blocks, LCBs). LCBs were aggregated to the gene level using their start and end coordinates. If the start or end coordinate of an LCB fell into a known gene, it was assigned to the gene. In cases where multiple LCBs were assigned to the same gene, the minimum and maximum coordinates of these LCBs became the start and end coordinates, respectively, for the gene-level loci. We manually filtered the obtained LCBs to identify regions that are shared among the strains that showed antimicrobial activities but are absent among the strains that did not show the activity. Annotations were added to the resulting file using the GTF file of *P. protegens* PBL3 obtained from NCBI and supplemented by KEGG functional annotations from anvi’o. Operons in the *P. protegens* PBL3 genome were predicted using the PathoLogic program of Pathway Tools v29.0 (Peter D. Karp, 2024).

### Phylogenetic analysis and Average Nucleotide Identity (ANI)

The phylogenomic analysis on *P. protegens* PBL3 and the six genomes followed the “Pangenomic + Phylogenomics” workflow (Delmont & Eren, 2018). Single-copy core gene clusters were extracted from the pangenomics profile and used in the phylogenomic analysis. Anvi’o’s anvi-gen-phylogenomic-tree utilizes FastTree (Price et al., 2010) by default to generate a phylogenomic tree. Average nucleotide identity (ANI) for the seven genomes was computed using the anvi’o command anvi-compute-genome-similarity (Pritchard et al., 2016; Richter & Rosselló-Móra, 2009).

## Results and Discussions

### Genetically related *Pseudomonas* spp. strains exhibit differential antimicrobial activity against *B. glumae*

To facilitate comparative genomic analysis for gene mining in *P. protegens* PBL3 we necessitated the identification of closely related *Pseudomonas* strains with fully sequenced genomes and contrasting phenotypes. For that purpose, we obtained six *Pseudomonas* spp. strains: *P. fluorescens* 513, *P. fluorescens* 1550, *P. protegens* 3295, *P. fluorescens* 1098, *P. fluorescens* 4488, and *P. putida* 4512 and extracted their respective secretomes to evaluate their antimicrobial activity against *B. glumae* and compare it with the antimicrobial activity of *P. protegens* PBL3. The results showed that *B. glumae* grown in KB medium alone (control) reached an OD_600_ of 2.5, whereas the growth of *B. glumae* in KB supplemented with the *P. protegens* PBL3 secretome was reduced by 88% as previously reported (Dahal et al., 2024). Among the six additional *Pseudomonas* strains tested, the secretomes of *P. fluorescens* 513, *P. fluorescens* 1550, and *P. protegens* 3295 exhibited antimicrobial activity against *B. glumae* similar to the antimicrobial activity observed with *P. protegens* PBL3, causing a reduction in *B. glumae* growth by 87%, 81%, and 87%, respectively. In contrast, the secretomes of *P. fluorescens* 1098, *P. fluorescens* 4488, and *P. putida* 4512 had no antimicrobial activity against *B. glumae,* and the levels of *B. glumae* growth were equivalent to those under control conditions (Figure 1A). Thus, *P. fluorescens* 513, *P. fluorescens* 1550, and *P. protegens* 3295 were considered as *B. glumae* antimicrobial-positive strains, whereas *P. fluorescens* 1098, *P. fluorescens* 4488, and *P. putida* 4512 were considered as *B. glumae* antimicrobial-negative strains.

**Figure. 1.**
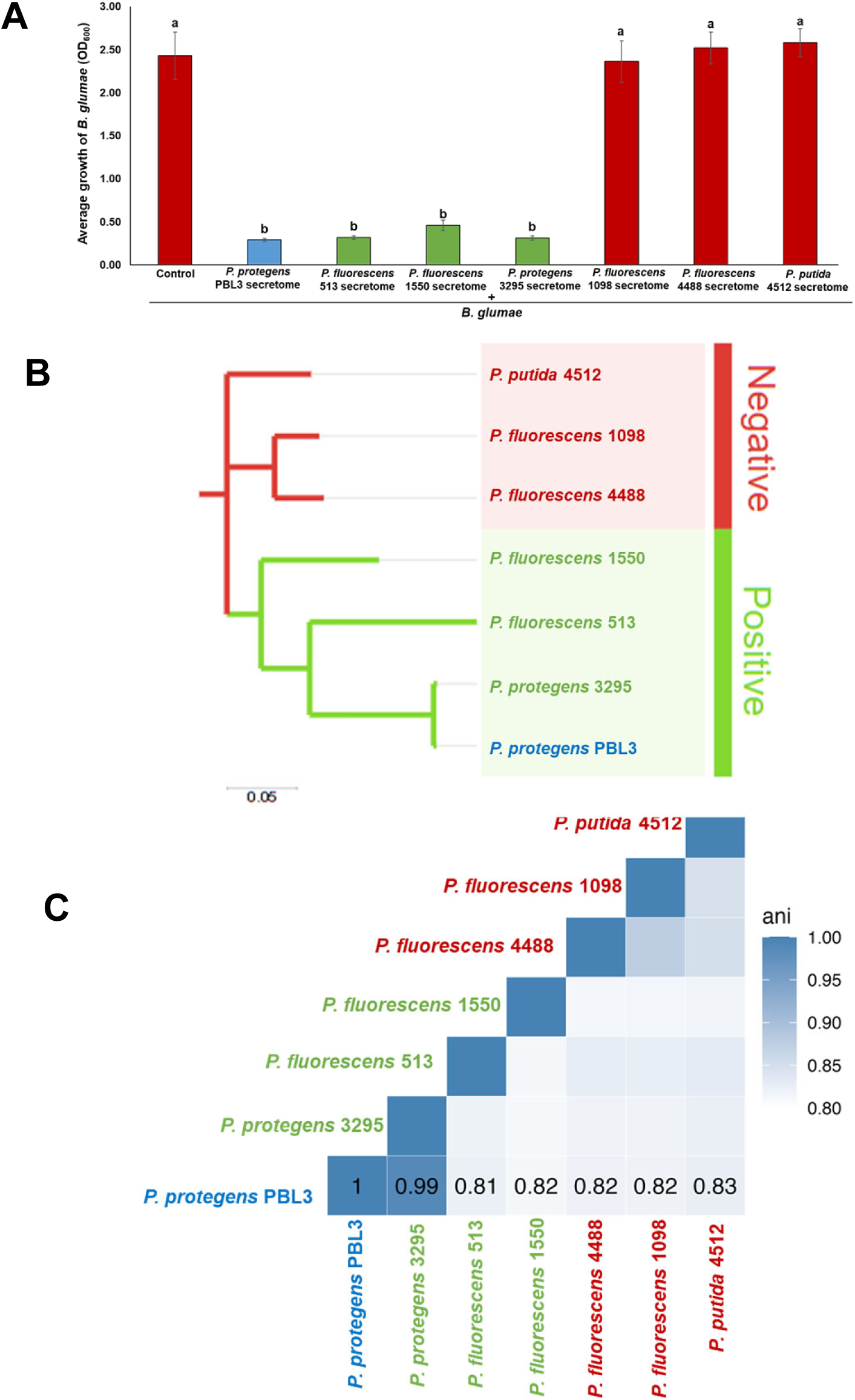
Antimicrobial activity of different strains of *Pseudomonas* against *B. glumae* and their genomic relatedness. **(A)** *B. glumae* at an optical density at 600 nm (OD_600_) of 0.2 was diluted in KB broth and mixed with water (control), and secretome from different strains of *P. protegens* PBL3 (treatments). The growth of *B. glumae* was evaluated after 18h. *Pseudomonas* spp. strains lacking antimicrobial activity are shown in red, *Pseudomonas* spp. strains with antimicrobial activity in green, and *P. protegens* PBL3 in blue. Bars represent the average growth of *B. glumae* with error bars indicating standard deviations from three biological replicates in three independent experiments. Statistical significance indicated by letters above the bars was determined using a one-way analysis of variance with Tukey’s significant difference test (P value < 0.05). **(B)** Phylogenomic tree of seven *Pseudomonas* strains inferred from single-copy core genes using anvi’o. Branch lengths are proportional to the number of substitutions per site, as indicated by the scale bar. Strains are grouped based on antimicrobial phenotype, with antimicrobial-negative strains indicated in red, antimicrobial-positive strains in green, and *P. protegens* PBL3 highlighted in blue. **(C)** Pairwise Average Nucleotide Identity (ANI) heatmap showing genomic relatedness of *P. protegens* PBL3 with the other *Pseudomonas* strains, calculated using anvi’o. Color intensity reflects ANI values, with darker blue indicating higher nucleotide identity. Antimicrobial phenotype coloring is consistent with panel B.

To assess the evolutionary relationship of the selected six *Pseudomonas* strains and *P. protegens* PBL3, we generated a phylogenomic tree and found that the three strains with antimicrobial activity against *B. glumae* (*B. glumae* antimicrobial-positive strains): *P. protegens* 3295, *P. fluorescens* 513, and *P. fluorescens* 1550 are in the same cluster with *P. protegens* PBL3, whereas the strains lacking antimicrobial activity against *B. glumae* (*B. glumae* antimicrobial-negative strains): *P. fluorescens* 4488, *P. fluorescens* 1098, and *P. putida* 4512 were in a different cluster (Figure 1B). Genome relatedness between *P. protegens* PBL3 and the other strains was further assessed using Average Nucleotide Identity (ANI), which showed 99% identity between *P. protegens* PBL3 and *P. protegens* 3295, while all other strains, regardless of clade position, shared ∼81–83% identity with *P. protegens* PBL3 (Figure 1C). This indicates that the selected strains are genetically related to *P. protegens* PBL3, ensuring that any differences identified in subsequent pan-genomic comparisons are more likely linked to antimicrobial activity rather than unrelated genetic divergence.

### *Pseudomonas* strains harbor distinct, strain-specific repertoires of biosynthetic gene clusters (BGCs)

Before starting genome comparisons, we curated the draft genomes of *Pseudomonas* strains *P. fluorescens* 513, *P. fluorescens* 1550, *P. protegens* 3295, *P. fluorescens* 1098, *P. fluorescens* 4488, and *P. putida* 4512 by scaffolding them against the *P. protegens* PBL3 reference genome. Scaffolding reduced the number of contigs while increasing the length of contigs, as demonstrated by higher N50 values and obtaining an L50 of one for all of the six *Pseudomonas* strains (Supplementary Table 1). L50 of one indicated that a single contig was sufficient to cover half of the genome assembly. This curation provided a fewer, longer contigs resulting in a more complete and less fragmented genome assembly.

We previously found that *P. protegens* PBL3 encoded 12 distinct BGCs predicted to encode secondary metabolites (Ortega et al., 2020). To gain insight into the coding capacity of the additional six *Pseudomonas* spp. with differential antimicrobial activity against *B. glumae*, we compared their BGC profiles against the *P. protegens* PBL3 BGC profile using antiSMASH v8.0.4. (Figure 2). This analysis revealed *P. protegens* PBL3 predicted to encode four new BGCs: protegencin, hydrogen cyanide, lankacidin C, and terpene-precursor. Comparative analysis of BGCs across all *Pseudomonas* strains with and without antimicrobial activity against *B. glumae* identified six BGCs predicted to encode arylpolyene APE Vf, fengycin, Pf-5 pyoverdine, N-acetylglutaminylglutamine amide, lankacidin C, and a terpene-precursor; BGC predicted to encode hydrogen cyanide was present in six of the strains and absent in *P. putida* 4512. When comparing *B. glumae* antimicrobial-positive strains, *P. protegens* 3295, and *P. protegens* PBL3, which have the closest genetic relationship, have seven common BGCs: 2,4-DAPG, protengencin, orfamide A/orfamide C, cydlodipeptides, enantio-pyochelin, pyrrolnitrin, lipopeptide 8D1-1/8D1-2. In contrast to *P. protegens* PBL3, *P. protegens* 3295 lacks pyoluteorin and bacteriocin BGCs and includes two additional BGCs predicted to encode alginate and hserlactone. The additional *B. glumae* antimicrobial-positive strains within the *P. fluorescens* group have genomes that predict specific BGC, such as viscosin and pyochelin in *P. fluorescens* 513; thanafactin A and histicorrugatin in *P. fluorescens* 1550; both *P. fluorescens* 513 and *P. fluorescens* 1550 genomes contain BGCs predicted to encode fragin, which is also encoded by the *B. glumae* antimicrobial-negative strain, *P. fluorescens* 4488. Additionally, *P. fluorescens* 1550 encoded 2,4-diacetylphloroglucinol, which is shared with both *P. protegens* strains, as well as butyrolactone and lanthipeptide class-ii cluster, which were also present in the *B. glumae* antimicrobial-negative strain, *P. fluorescens* 4488. Further, *Pseudomonas* spp. strains lacking antimicrobial activity against *B. glumae* encoded additional BGCs: lokisin in *P. fluorescens* 4488 and *P. fluorescens* 1098; nematophin, chitinimide B, and an NRPS-independent siderophore in *P. fluorescens* 1098; and a phenazine BGC in *P. putida* 4512.

**Figure 2.**
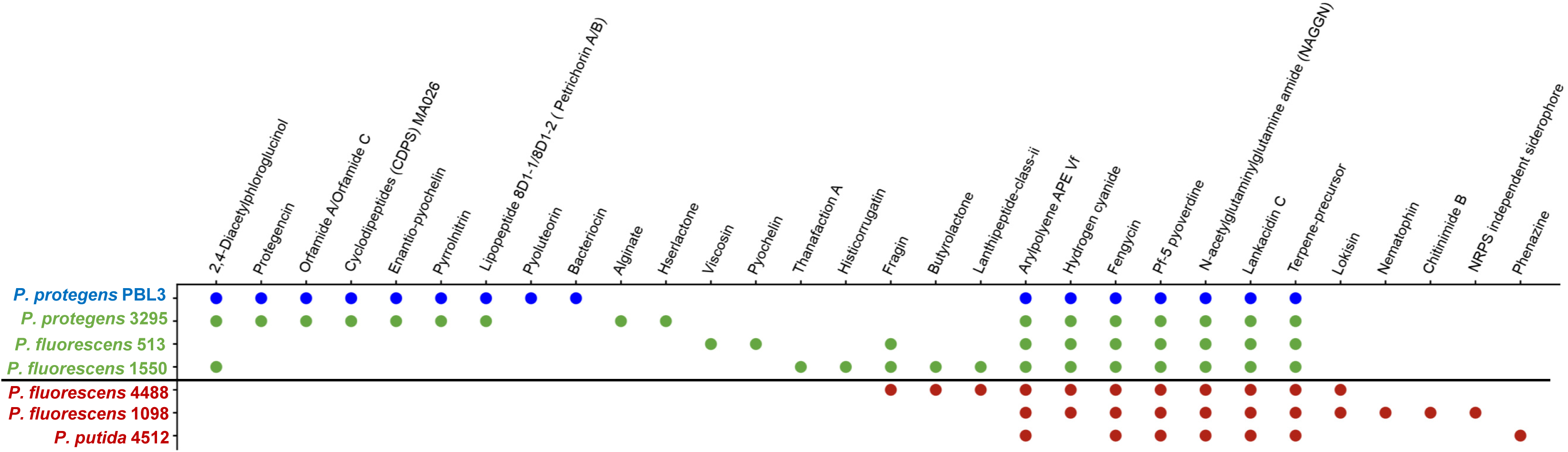
Distribution of predicted biosynthetic gene clusters (BGCs) across *Pseudomonas* strains with differential antimicrobial activity against *B. glumae*. Presence-absence matrix of BGC predicted using antiSMASH v8.0.4 across seven *Pseudomonas* genomes. Each column represents a predicted BGC, annotated by the putative metabolite class or product, and each row represents an individual strain (colored by phenotype: *P. protegens* PBL3 in blue, antimicrobial-positive in green, and antimicrobial-negative in red). Colored circles indicate the presence of a given BGC in a strain. The horizontal line separates antimicrobial-positive and antimicrobial-negative strains.

Overall, this analysis revealed that *Pseudomonas* spp. genomes with antimicrobial activity against *B. glumae* are endowed with more BGCs than those without it. Importantly, it appears that this antimicrobial activity is associated with combinations of BGCs rather than specific BGCs, as we initially hypothesized (Dahal et al., 2024). Although these results are insightful, this analysis using antismash did not allow us to define the specific BGCs associated with the antimicrobial activity of *P. protegens* PBL3. This may be due to antismash limitations regarding its dependency on existing rule-based detection, predicting canonical BGCs, and not capturing non-BGC genes or divergent biosynthetic pathways, such as regulatory, transport, secretion, or accessory metabolic genes that may contribute to antimicrobial activity (Kang et al., 2025).

### Comparative genomics identifies genomic regions uniquely associated with antimicrobial activity in *P. protegens* PBL3

The comparative BGCs profiling analysis using antismash challenged our hypothesis regarding the conservation of BGCs among strains with activity and the absence of those BGCs in the strains without activity. Thus, we opted to generate more robust information on genomic regions associated with antimicrobial activity by comparative genomics. We initially compared all seven genomes at the amino acid level using anvi’o to identify core gene clusters defined as an orthologous group of homologous genes identified across genomes in the anvi’o pangenome analysis and not necessarily physical linkage. The pangenome of the 7 strains included 40,762 genes, of which 22,518 genes (55.0%) formed the consensus core genome (Figure 3, dark blue band). Within phenotypic groups, strains with antimicrobial activity (*B. glumae* antimicrobial-positive) (including *P. protegens* PBL3) shared 1,684 core genes (310 clusters) (Figure 3, orange band), while strains lacking antimicrobial activity (*B. glumae* antimicrobial-negative) shared 4008 core genes (778 clusters) (Figure 3, purple band). To identify genomic regions potentially associated with antimicrobial activity, we focused on genes within the clusters present in all *B. glumae* antimicrobial-positive strains and *P. protegens* PBL3 but absent in all *B. glumae* antimicrobial-negative strains (Figure 3, orange band with star). This analysis revealed 98 genes (96 clusters) unique to strains with antimicrobial activity, as genomic differences associated with antimicrobial activity (Figure 3, Supplementary Table 2). These accessory 98 genes represent potential candidates for functional characterization.

**Figure 3.**
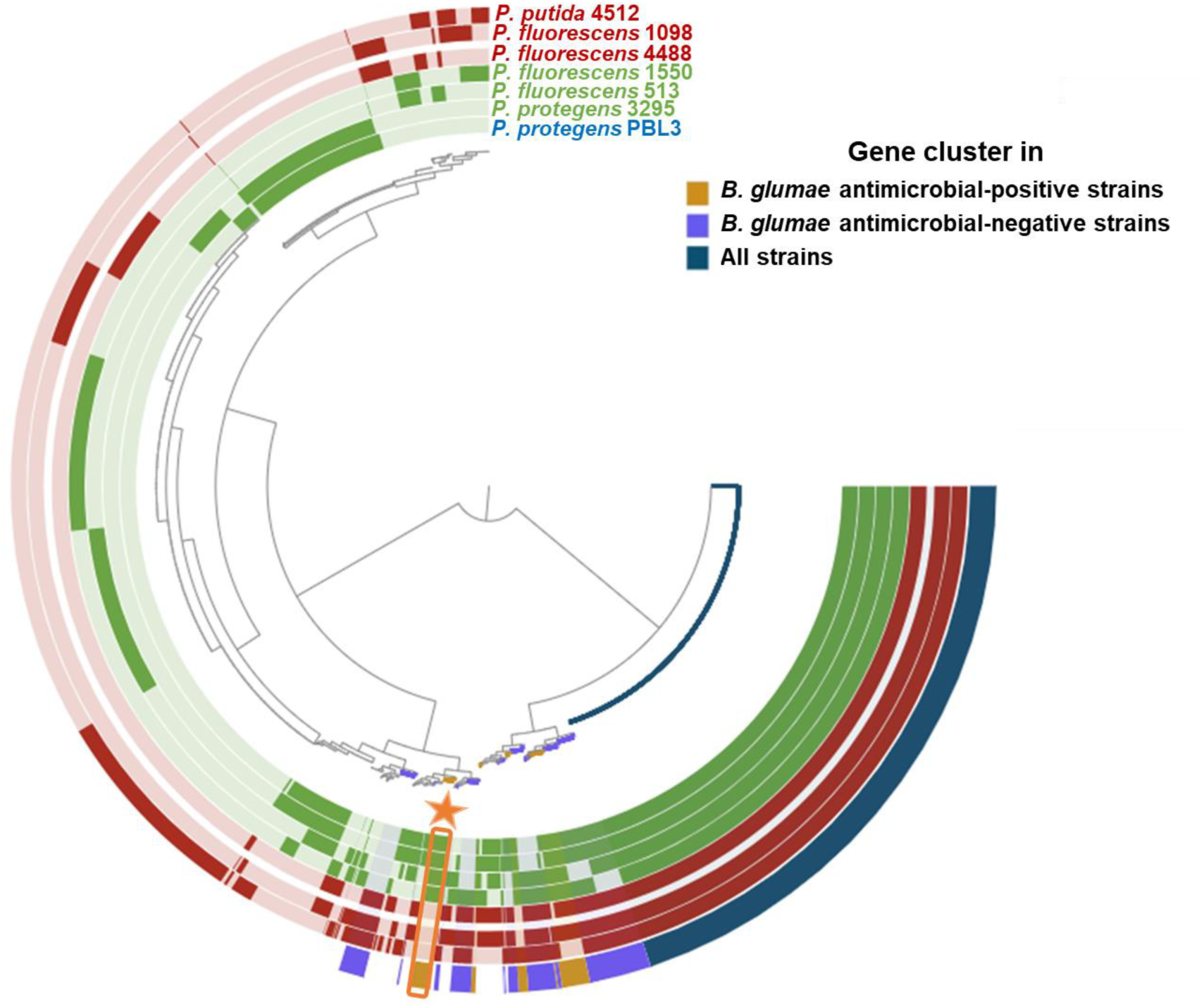
Comparative genomics identifies genes uniquely associated with antimicrobial activity against *B. glumae.* Circular pangenome visualization of seven *Pseudomonas* genomes generated using the anvi’o pangenomics workflow. The circular dendrogram was constructed based on the presence or absence of gene clusters (orthologous gene families) across genomes. Concentric rings represent individual genomes for *B. glumae* antimicrobial-positive strains in green, *B. glumae* antimicrobial-negative strains in red; colored blocks indicate presence of gene clusters in that genome. The outermost ring summarizes gene cluster distribution patterns across phenotypic groups: orange blocks indicate gene clusters exclusive to antimicrobial-positive strains, purple blocks indicate gene clusters exclusive to antimicrobial-negative strains, and dark blue blocks denote gene clusters present in both antimicrobial-positive and antimicrobial-negative strains. The orange star highlights a genomic region enriched in gene clusters uniquely conserved among antimicrobial-positive strains but absent from antimicrobial-negative strains.

Since anvi’o pan-genome analysis does not capture subtle differences at the nucleotide level, such as small insertions, deletions, or point mutations, we complemented the anvi’o analysis with a whole-genome multiple sequence alignment using progressiveMauve to generate more robust information. progressiveMauve identified 223 unique genomic hits that are present in *P. protegens* PBL3 and in other strains showing antimicrobial activity (*B. glumae* antimicrobial-positive strains) but absent in strains lacking antimicrobial activity (*B. glumae* antimicrobial-negative strains) (Supplementary Table 3). Out of these, 161 correspond to annotated genes, while 62 regions are uncharacterized. The uncharacterized regions suggested that genes encode novel mechanisms that contribute to antimicrobial activities.

Together, these two complementary approaches resulted in 188 annotated non-redundant putative genes potentially associated with antimicrobial activity, 27 genes unique to anvi’o and 87 genes unique to progressiveMauve, with 74 genes overlapping between the two analyses (Supplementary Table 4).

### Antimicrobial-associated genes localize within predicted biosynthetic gene clusters (BGCs) in *P. protegens* PBL3

To determine whether any of the 188 genes uniquely present in *P. protegens* PBL3 and the *B. glumae* antimicrobial-positive strains were directly associated with known secondary metabolite biosynthesis, we examined their genomic locations relative to the BGCs predicted in *P. protegens* PBL3 using antismash (Figure 2). This mapping revealed that seven of the 188 candidate genes fell within six BGCs previously identified (Figure 4). Genes HH203_RS10555, HH203_RS26235 and HH203_RS19805 mapped to orfamide A/orfamide C cluster, terpene-precursor cluster and pyoverdine cluster, respectively and encode core biosynthetic genes. Two additional genes, HH203_RS19700 mapped to the pyoverdine cluster and HH203_RS22555 mapped to the lipopeptide 8D1-1/8D1-2 (Petrichorin A/B) cluster and are associated with additional biosynthetic functions. Within the 2,4-DAPG cluster, gene HH203_28595 encodes an RND-family efflux transporter, indicating that antimicrobial activity may also depend on metabolite export associated with BGCs function. The remaining candidate, HH203_RS12645 in the hydrogen cyanide cluster, encodes a YdcA family protein, whose function remains poorly characterized. Although we had already identified these clusters with the *P. protegens* PBL3 genomic sequencing, these results highlight specific genes within those clusters as uniquely present in *B. glumae* antimicrobial-positive strains, suggesting that those genes are essential for the production of a specific metabolite different from the commercial standards used in previous analyses (Dahal et al., 2024). This pattern is consistent with biosynthetic processes mediated by the activities of non-ribosomal peptide synthetases (NRPS), such as the synthesis of orfamide, pyoverdine and lipopeptides; and polyketide synthases, such as the synthesis of 2,4-DAPG whose activities can generate a variety of structurally diverse molecules without changes at the genomic level (Gross & Loper, 2009; Hertweck, 2009; Höfte, 2021; Payne et al., 2016).

**Figure 4.**
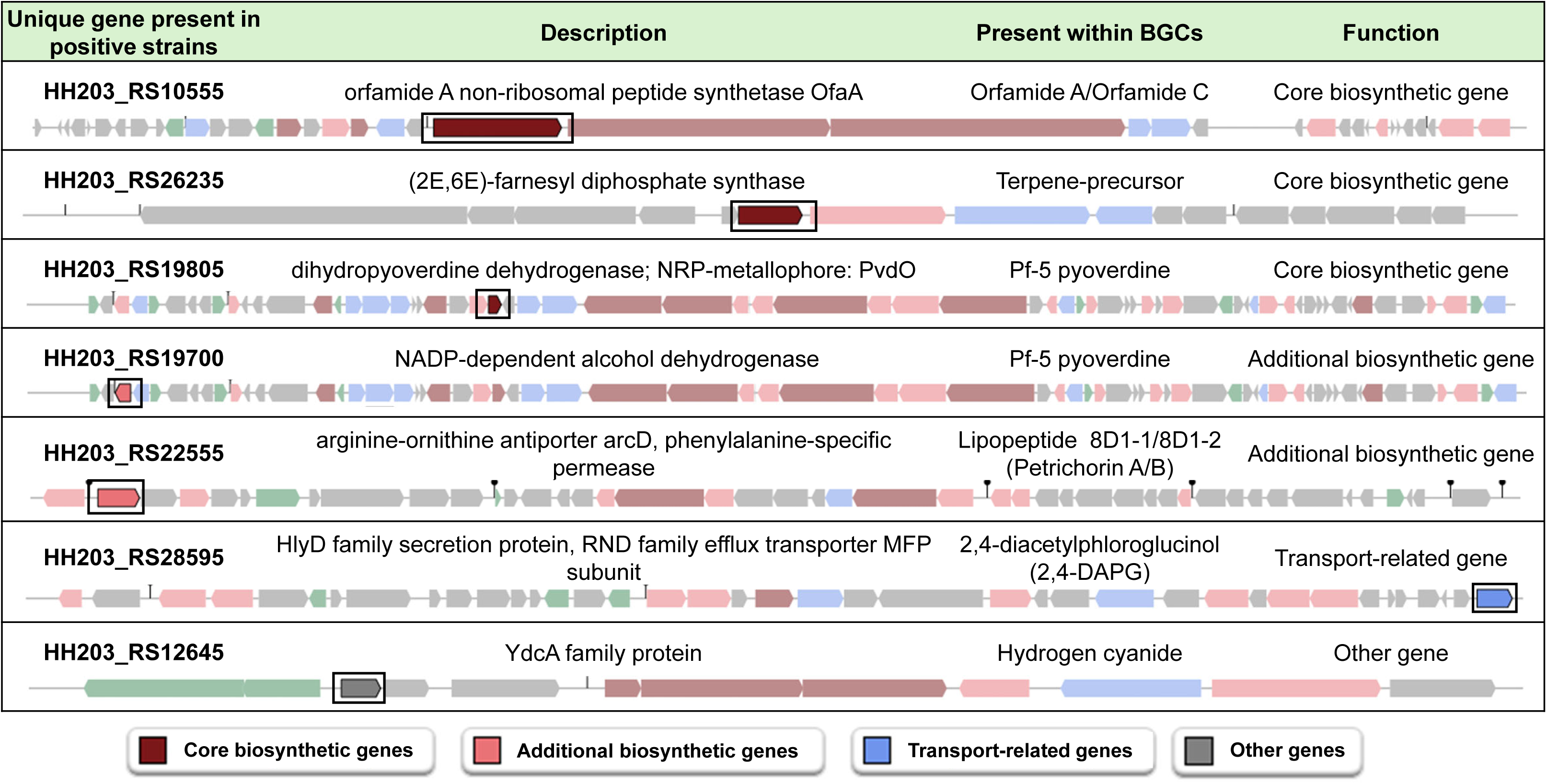
Antimicrobial-associated genes mapped to predicted biosynthetic gene clusters (BGCs) in *P. protegens* PBL3. Linear genomic representations of biosynthetic gene clusters (BGCs): orfamide A/C, terpene precursor, Pf-5 pyoverdine, lipopeptide 8D1-1/8D1-2 (Petrichorin A/B), 2,4-DAPG, and HCN in the *P. protegens* PBL3 genome predicted by antiSMASH. Each block arrow represents a gene and its orientation within the cluster. Genes uniquely conserved among antimicrobial-positive strains are highlighted with black outlines. Genes are color-coded by functional category: core biosynthetic genes (maroon), additional biosynthetic genes (light red), transport related genes (blue), and other genes (gray).

### KEGG-based functional categorization reveals a multi-functional network of antimicrobial-associated genes in *P. protegens* PBL3

The seven genes that are located in previously known metabolite-producing clusters represent only a small fraction of the 188 candidate genes identified through comparative genomics. This suggests that antimicrobial activity-associated genes in *P. protegens* PBL3 extend well beyond classical secondary metabolite pathways. To better understand the potential biological roles of the candidate genes, we performed KEGG-based functional annotation of all 188 genes uniquely present in *P. protegens* PBL3. KEGG Orthology (KO) accessions assigned by anvi’o were mapped to the KEGG Pathway and BRITE databases, followed by manual curation where necessary to generate functionally meaningful groupings. This analysis enabled us to categorize the genes into major functional classes, including metabolism, enzyme class, genetic information processing, environmental information processing, and cellular processes (Figure 5, Supplementary Table 5), providing a broader view of the molecular functions potentially contributing to antimicrobial activity. To capture the complete repertoire of candidates, KEGG analysis was performed separately for high-confidence genes detected by both comparative methods (Figure 5, light purple bars) and for those identified uniquely by either anvi’o (Figure 5, light brown bars) or progressiveMauve (Figure 5, light blue bars).

**Figure 5.**
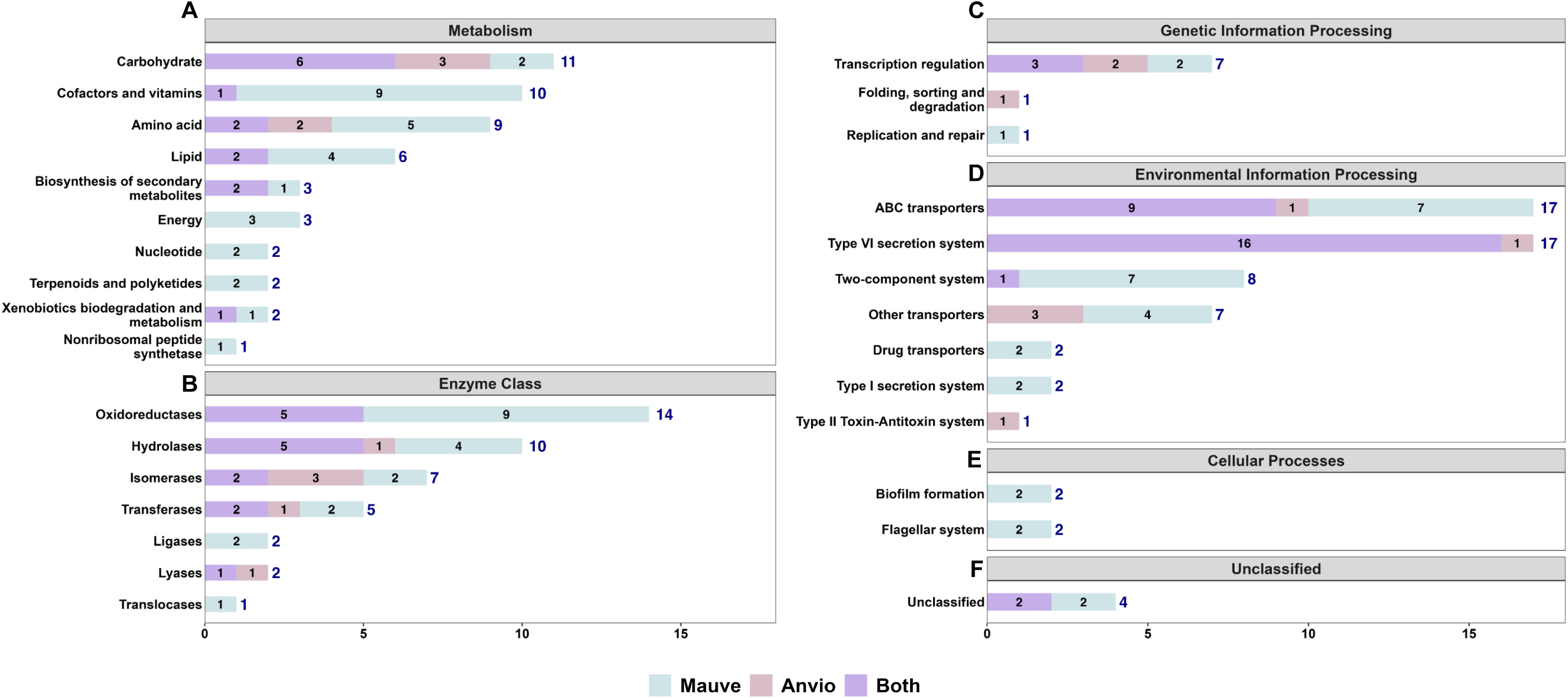
KEGG-based functional classification of genes uniquely associated with antimicrobial activity in *P. protegens* PBL3. Stacked bar plots summarizing KEGG functional assignments for genes uniquely present in antimicrobial-positive strains (including *P. protegens* PBL3) and absent in antimicrobial-negative strains. Genes were annotated using KEGG Orthology (KO) assignments and grouped into major functional categories indicated in panel as **(A)** Metabolism, **(B)** Enzyme class, **(C)** Genetic information processing, **(D)** Environmental information processing, **(E)** Cellular processes, and **(F)** Unclassified. Bars represent the number of genes assigned to each group within the major functional category, with color coding indicating the method by which genes were identified: progressiveMauve only (light blue), anvi’o only (light brown), or overlapping between both methods (light purple). Numbers at the end of each bar indicate total gene counts per functional group.

Because a single gene can function in multiple pathways, some genes were represented across both metabolic and enzyme class category (Supplementary Table 5). Within the metabolism network, genes were distributed across ten functional groups involved in the biosynthesis of structurally diverse antimicrobial metabolites. The largest representation was in carbohydrate metabolism, followed by metabolism of cofactors and vitamins, amino acid metabolism, lipid metabolism, biosynthesis of secondary metabolites, energy metabolism, nucleotide metabolism, terpenoids and polyketides metabolism, xenobiotic biodegradation and metabolism and nonribosomal peptide synthetase metabolism (Figure 5A). These metabolic pathways provide the essential precursors, biochemical intermediates, and cofactors that support antimicrobial metabolite biosynthesis in *P. protegens* PBL3. Enzyme classification further resolved the repertoire of catalytic functions encoded by these genes. The distribution was dominated by oxidoreductases, followed by hydrolases, isomerases, transferases, ligases, lyases, and translocases (Figure 5B). Many of these enzymes mediate activation, modification, and diversification reactions that enhance the structural complexity and bioactivity of antimicrobial compounds. Collectively, these non-exclusive groups define the biochemical versatility and broad metabolic reservoir supporting antimicrobial metabolite production in *P. protegens* PBL3.

Beyond metabolism and enzymatic functions, KEGG annotation revealed genes uniquely associated with genetic information processing, environmental information processing, and cellular processes (Supplementary Table 5), which collectively contribute to regulation, secretion, transport, and stress adaptation associated with antimicrobial activity. A total of nine genes were classified under genetic information processing, including seven transcription regulation and one gene each for protein folding, sorting, and degradation; and replication and repair (Figure 5C). Fifty-four genes were classified under environmental information processing, encompassing diverse transport systems, secretion machinery, and regulatory modules. The largest groups included ABC transporters and the type VI secretion system, each with seventeen genes. Eight genes encoded two-component systems, seven genes corresponded to other transporters, two genes to drug transporters, two genes to the Type I secretion system, and one gene to a Type II toxin–antitoxin system (Figure 5D). Four genes were classified under cellular processes, including two involved in biofilm formation and two in the flagellar system (Figure 5E). Four genes lacked definitive pathway assignments and were categorized as unclassified (Figure 5F).

Collectively, genes classified under genetic information processing, environmental information processing, and cellular processes form an integrated regulatory and export network that complements core metabolic and enzymatic pathways. The KEGG-based functional classification thus highlights a complex, multi-functional network of candidate genes in antimicrobial-producing strains, encompassing biosynthetic enzymes, transporters, secretion systems, and transcriptional regulators. These findings suggest that *P. protegens* PBL3 employs not only classical biosynthetic pathways but also coordinated regulatory, metabolic, and export modules to synthesize and deploy antimicrobial compounds.

### Identification of antimicrobial-associated gene clusters in *P. protegens* PBL3 genome uncovers multiple functional categories

To contextualize the 188 candidate genes uniquely present in *P. protegens* PBL3 and the antimicrobial-positive strains, we examined their physical distribution across the genome. Manual inspection using Integrative Genomic Viewer (IGV) revealed that these genes were not randomly distributed but formed 25 discrete genomic clusters ranging in size from 1.2 Kb to 34.7 kb that were numbered based on their genomic position in *P. protegens* PBL3 (Figure 6). Each of these genomic clusters comprises sets of adjacent genes encompassing two to twenty-three genes separated by no more than 1 kb and frequently containing one or more putative operons. Because specialized metabolic functions and antimicrobial pathways in *Pseudomonas* are often encoded in operon-like loci, this architecture suggests that these regions may represent candidate antimicrobial-associated modules. Functional annotation of the genes based on gene description within each cluster exhibits a coherent biological theme, allowing us to infer higher-order functional modules for each cluster (Table 2, Supplementary Table 6).

**Figure 6.**
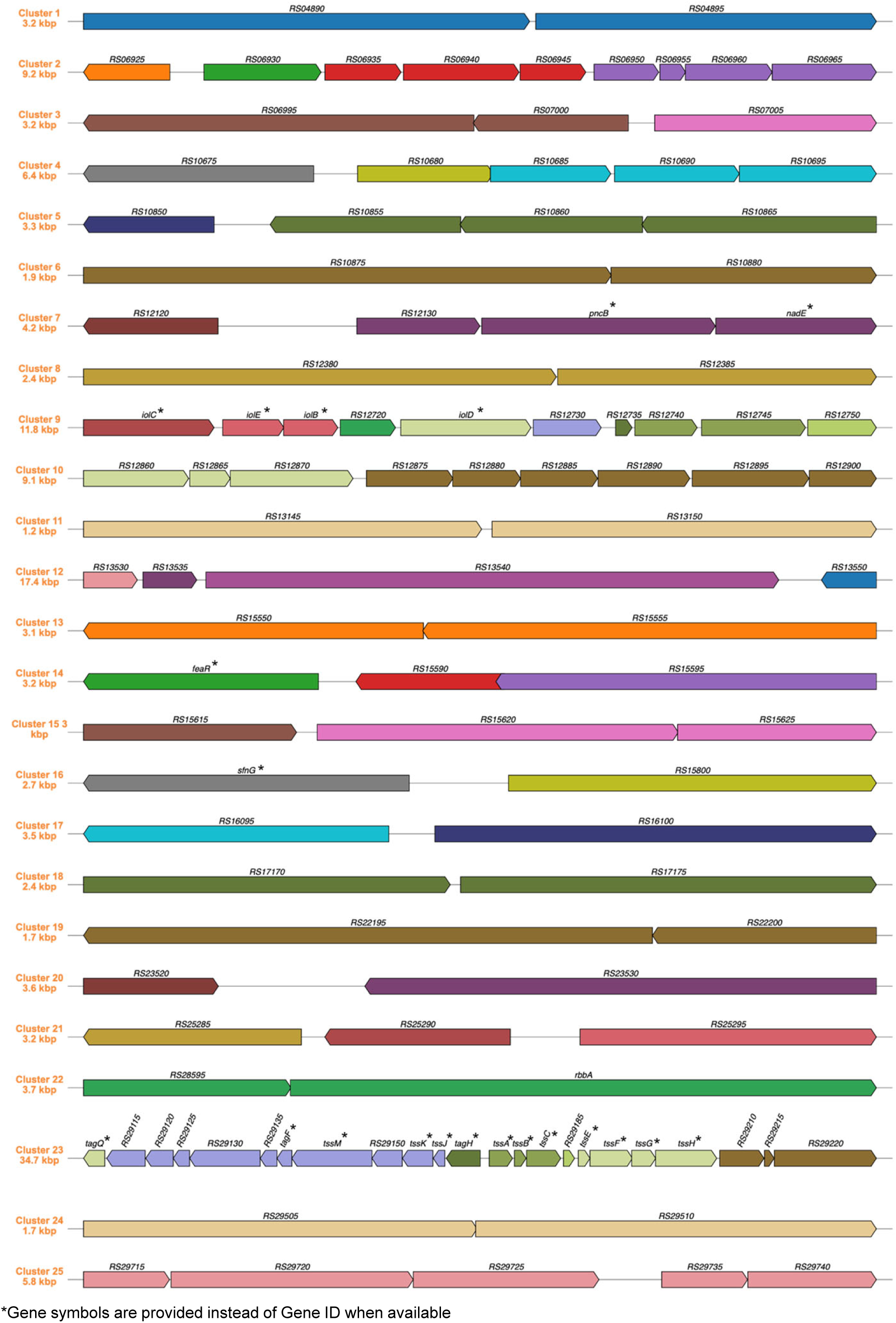
Genomic organization of antimicrobial-associated gene uniquely present in *P. protegens* PBL3. Linear maps of 25 discrete genomic clusters identified in *P. protegens* PBL3 that harbor genes uniquely present in antimicrobial-producing strains and absent in antimicrobial-negative strains. Clusters were defined as sets of adjacent genes separated by ≤1 kb and frequently containing one or more putative operons. Arrows represent predicted open reading frames oriented by transcriptional direction; genes sharing the same color within a cluster indicate putative operons. Clusters are numbered (Clusters 1-25) according to their genomic position in the *P. protegens* PBL3 genome, with total cluster sizes indicated in kilobases (kbp).

**Table 2.**
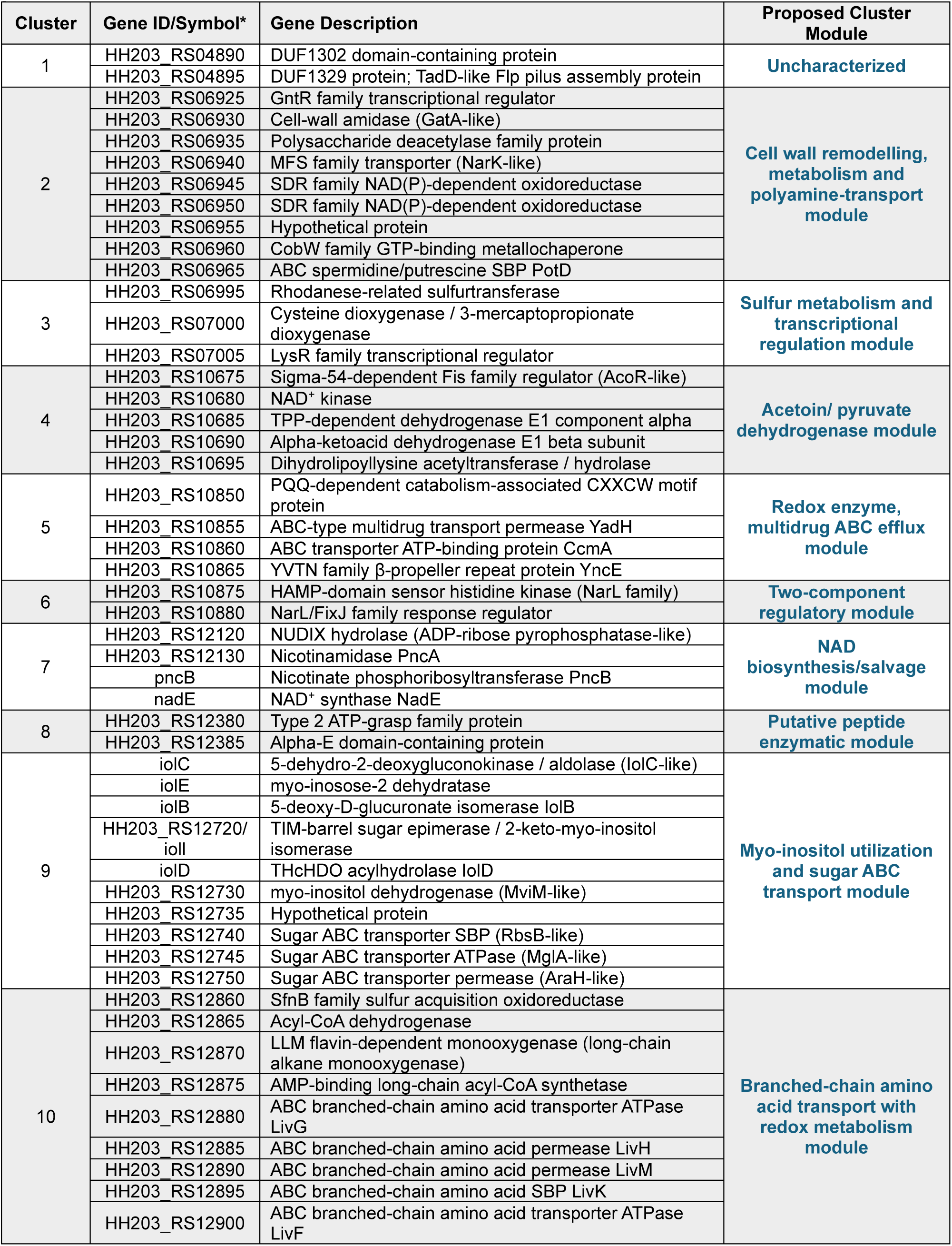
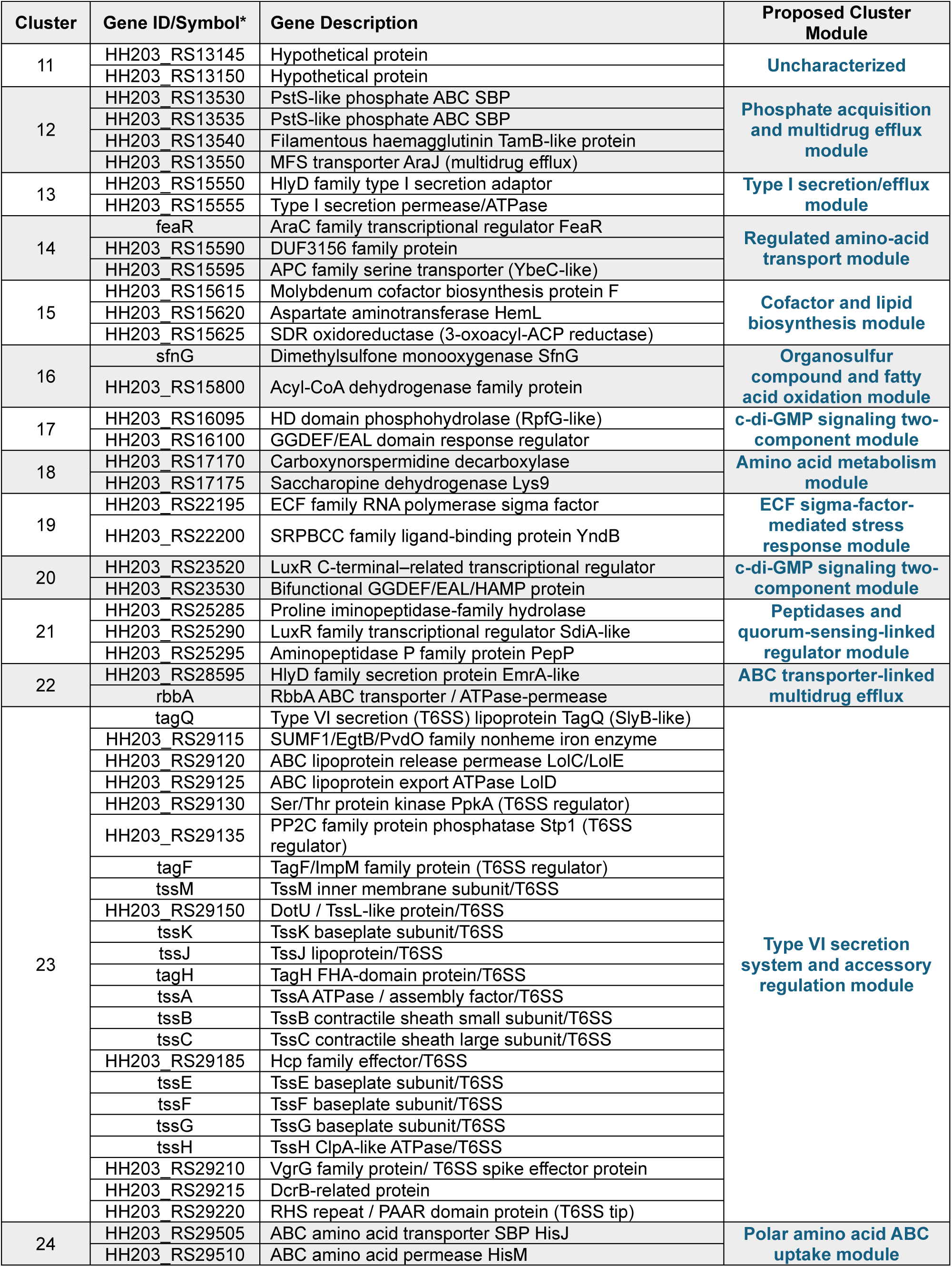
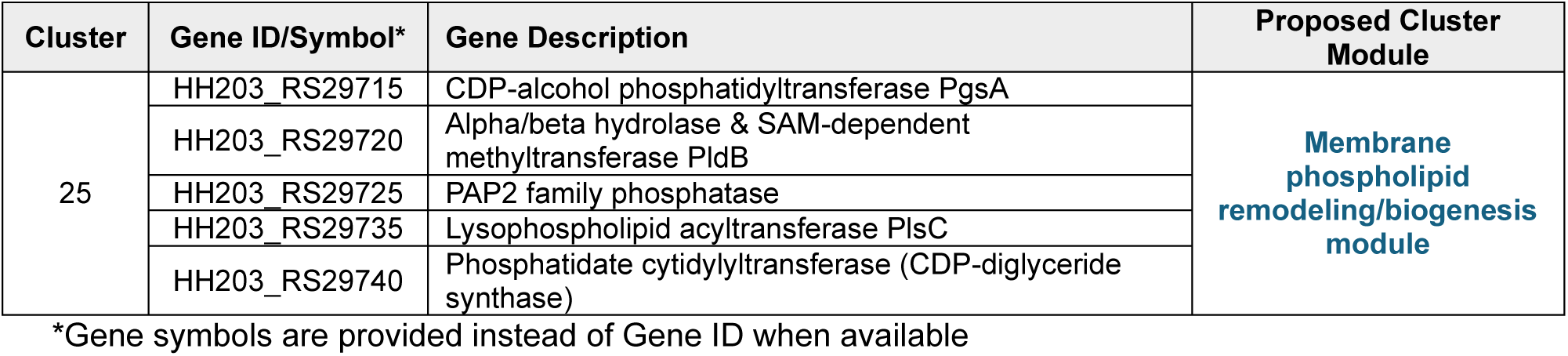
Antimicrobial-associated genes in *P. protegens* PBL3 are organized into 25 putative functional clusters.

Several clusters encode metabolic or biosynthetic modules, including sulfur- and amino-acid metabolism (Clusters 3 and 18), the NAD biosynthesis/salvage pathway (Cluster 7), acetoin/pyruvate metabolism (cluster 4), a cofactor and lipid biosynthesis module (Cluster 15), organosulfur and fatty-acid oxidation (cluster 16), and myo-inositol catabolism and uptake (Cluster 9). Within cluster 9 an *iol* locus encodes for myo-inositol catabolism (iolC/E/B/I/D, an inositol dehydrogenase, and ABC-type sugar transporters). Previous work in *Pseudomonas* has linked the *iol* locus to inositol-responsive behaviors, including fluorescent siderophore accumulation and competitive competence (Sánchez-Gil et al., 2023). In our assays, all *Pseudomonas* strains grew in myo-inositol medium even strains lacking the cluster (i.e. antimicrobial-negative strains), indicating that this locus is not necessarily required for myo-inositol catabolism but might influence production or accumulation of antimicrobial molecules.

Transport and efflux systems were also prominent and included a cell-wall remodeling and polyamine-transport module (Cluster 2), phosphate acquisition coupled to multidrug efflux (Cluster 12), ABC-linked multidrug secretion systems (Clusters 5 and 22), several amino-acid and sugar ABC transporters (Clusters 9, 14, and 24), branched-chain amino-acid transport with alkane/sulfonate redox metabolism (Cluster 10), and a dedicated Type I secretion system (Cluster 13). Cluster 19 represents an ECF sigma-factor mediated stress-response module. Cluster 21 links two peptidases with a LuxR-family regulator, suggesting peptide processing coupled to quorum-sensing–like regulatory control. Cluster 25 encodes key enzymes involved in membrane phospholipid biogenesis and remodeling, supporting membrane integrity. We also identified a subset of genomic regions (Clusters 1, 8, and 11) that remain largely uncharacterized, consisting mainly of DUF-containing or conserved hypothetical proteins without clear functional annotation, and therefore represent promising regions for future functional dissection of novel antimicrobial determinants.

Beyond these clusters, we also identified two component systems (TCS) and c-di-GMP signaling systems (Clusters 6, 17, and 20) uniquely present in *B. glumae* antimicrobial-positive strains, supporting a role for signal transduction in regulating secretome-mediated antimicrobial activity. TCS act as environmental signal transducers that enable bacteria to adapt to changing conditions and are recognized as broad regulators of antibiotic and secondary metabolite biosynthesis, thereby influencing biocontrol traits in beneficial bacteria such as *Pseudomonas, Bacillus, and Streptomyces* (Cruz-Bautista et al., 2023; Gaffney et al., 1994; Yu et al., 2023). Consistent with this, our comparative genomic analysis identified clusters 6, 17, and 20 as enriched for TCS related signaling and regulatory genes and further revealed the response regulator HH203_RS27265 (*creC*) though located outside these clusters, represents additional TCS as exclusively present in antimicrobial-positive strains. Interestingly, a deletion of *creC* in the antagonistic bacterium *Pseudomonas taiwanensis*, reduces its antimicrobial activity against the rice pathogen *Xanthomonas oryzae* pv. *oryzae* (*Xoo)* (Chen et al., 2020).

Our most significant result was uncovering Type VI secretion system (T6SS) within cluster 23 exclusively present in antimicrobial-producing strains. Remarkably, we identified 23 T6SS-related genes, including those encoding structural proteins (TssA–TssM), baseplate components, Hcp, VgrG, a PAAR tip effector, lipoproteins, and associated kinases/phosphatases (Coulthurst, 2019). Its completeness and regulatory composition strongly support a potential role in interbacterial antagonism and delivery of antimicrobial molecules. Although T6SS is classically viewed as a contact-dependent interbacterial weapon (Navarro-Monserrat & Taylor, 2023), emerging research suggests broader roles in *Pseudomonas*, including links to the secretion and regulation of bioactive metabolites such as pyoverdine (Chen et al., 2016). Consistent with this, we identified two genes within the pyoverdine BGC uniquely present in antimicrobial-positive strains, supporting a potential functional coupling between pyoverdine production and T6SS-associated competitive fitness. One possible model is that T6SS contributes indirectly to secretome-mediated antimicrobial activity by shaping metabolic output, export, or regulatory states that favor the accumulation of toxic or competitive metabolites in the extracellular milieu. Such coordination would allow antimicrobial-positive strains to deploy a contact-independent chemical offense while retaining the capacity for T6SS-mediated competition in natural communities.

Collectively, the organization of these 25 clusters demonstrates that the genomic basis of *P. protegens* PBL3 antimicrobial activity is multifactorial, comprising coordinated metabolic, regulatory, secretion, and transport systems rather than a single biosynthetic pathway. Overall, our results highlight comparative genomics as a robust framework to identify genomic regions associated with the antimicrobial activity of *P. protegens* PBL3 against *B. glumae* to prioritize genes and pathways for future functional characterization.

## Acknowledgements

We thank Dr Daniel Schatmann’s lab (UNL) for providing the *Pseudomonas* spp. strains used in the study and JGI Integrated Microbial Genomes & Microbiomes (IMG/M) for the *Pseudomonas* spp. genome sequence data. This project was funded by the National Science Foundation (Award # 2334721) and the National Institute for Food and Agriculture (Award # 2024-67013-43202).

## Supplementary Tables

**Supplementary Table 1.** Genome assembly statistics of *Pseudomonas* strains before (A) and after (B) scaffolding, using the *P. protegens* PBL3 genome as reference.

**Supplementary Table 2.** Anvi’o pangenome workflow identified genes exclusively present in *P. protegens* PBL3 and antimicrobial-positive strains but absent in antimicrobial-negative strains.

**Supplementary Table 3.** Whole-genome multiple sequence alignment using progressiveMauve identified genes exclusively present in *P. protegens* PBL3 and antimicrobial-positive strains but absent in antimicrobial-negative strains.

**Supplementary Table 4.** Integration of comparative genomics approaches identified genomic regions uniquely associated with antimicrobial activity in *P. protegens* PBL3.

**Supplementary Table 5.** KEGG-based functional classification of candidate genes associated with antimicrobial activity.

**Supplementary Table 6.** Detailed annotation of antimicrobial-associated genes in *P. protegens* PBL3 organized into 25 putative functional clusters.

